# Uncoupling of synaptic loss from amyloid burden by an Alzheimer’s disease protective variant of PLCγ2

**DOI:** 10.1101/2023.09.22.558987

**Authors:** Ryan J. Bevan, Emily Maguire, Thomas Phillips, Elena Simonazzi, Marieta Vassileva, Julie Williams, Philip R. Taylor

## Abstract

A rare coding missense variant (rs72824905; P522R) in *PLCG2* decreases the risk of late-onset Alzheimer’s disease, but how this protective effect is mediated is unclear. Here we demonstrate a mechanism for this protection, the R522 variant of PLCγ2 alters microglial activity leading to a marked preservation of synaptic integrity and reduced peri-plaque microglial engulfment of synapses independently of amyloid burden. Our data advocate for a direct central role of PLCγ2 in mediating synaptic loss as part of the pathological process of Alzheimer’s disease (AD), prioritising it as a therapeutic target and modulator of disease.

---

A rare protective protein-coding missense variant in the *PLCG2* gene (rs72824905, P522R), which decreases the risk of developing late-onset Alzheimer’s disease (LOAD), was relatively recently identified^1^. Subsequent studies have confirmed association of the R522 variant with other dementias and increased longevity in centenarian studies, consolidating the evidence of a central role for *PLCG2* in neurodegenerative pathologies^2–4^.

The *PLCG2* gene encodes the enzyme Phospholipase C Gamma 2 (PLCγ2), an important regulatory hub gene involved in the hydrolysis of phosphatidylinositol (4,5) bisphosphate (PI(4,5)P2) into diacylglycerol (DAG) and inositol 1,4,5 trisphosphate (IP3), modulating intracellular Ca^2+^ and key protein kinase C signalling pathways^5^. In the brain, *PLCG2* is selectively expressed in microglia and is part of the same interaction network as *TREM2*, a significant susceptibility gene for AD^6,7^. Microglia, the major brain resident macrophages, are considered important for synaptic health, debris phagocytosis and inflammatory mediator production, and in AD are considered to play central roles in the pathology. Recently, we have seen the first successes in the development of therapeutic interventions for AD, with reagents that target and promote the removal of amyloid by microglia-driven pathways (e.g. Lecanemab, Donanemab)^8–10^. Such interventions modestly reduce cognitive decline in select patient subgroups but present with a notable risk of side effects, including brain swelling and microbleeds, necessitating the development of alternative and adjunctive therapeutic targets. Given the inherent druggability of PLCγ2, and its role in modulating macrophage function, understanding its direct involvement, if any, in the specific pathological processes, such as synapse loss, that are associated with diminishing cognition is crucial for its development as a therapeutic target.

In AD model mice, *Plcg2* is highly expressed in plaque-associated microglia, with its abundance increasing along the pathological timeline^11,12^. We previously demonstrated that the R522 variant of PLCγ2 is associated with hyperfunctionality, and results in marked changes in the endocytic and phagocytic capability of microglia and macrophages^13^. An initial study on a small cohort of male *Plcg2*^*R522*^ knock-in mice reported an increased number of P2ry12^+^ microglia in the cortex with no overt morphological differences compared to WT mice^14^. Thus, whilst we have observed clear impacts of the R522 variant on microglial function *in vitro*, its *in vivo* role in mediating microglial-specific protection from critical disease-relevant pathologies, such as progressive synapse loss, is unknown and would be potentially difficult to separate from any influence over plaque development.

To further these understandings, we initially analysed the impact of the P522R variants on microglial morphology and distribution *in vivo* (**Fig. 1**). Mice expressing *Plcg2*^*R522*^ at 6 months of age exhibited an increased density of Iba1^+^ microglial cell tiling in the dorsal hippocampus CA1 region (**Fig. 1 a-c**) and overlaying cortex (**Supp Fig. 1a-c**) when compared to *Plcg2*^*P522*^ expressing mice. There was no impact of sex on the manifestation of these *Plcg2* variant-related differences in microglial distribution. A morphological examination of the microglia in the hippocampus CA1 region revealed that the R522 variant was associated with diminished ramifications (**Fig 1 d-e**) and fewer terminal points (**Fig. 1 f**) than microglia with the P522 variant. We also noted a reduction in the overall microglia volume and an increase in the prevalence of lysosomal protein CD68, a commonly used marker of phagocytic activity, in *Plcg2*^*R522*^-expressing microglia compared to those expressing the P522 variant (**Fig. 1 g-i**). However, cell soma measurements reflecting volume and sphericity remained similar between the two genotypes (**Fig. 1 j-k**).

**Fig. 1:**
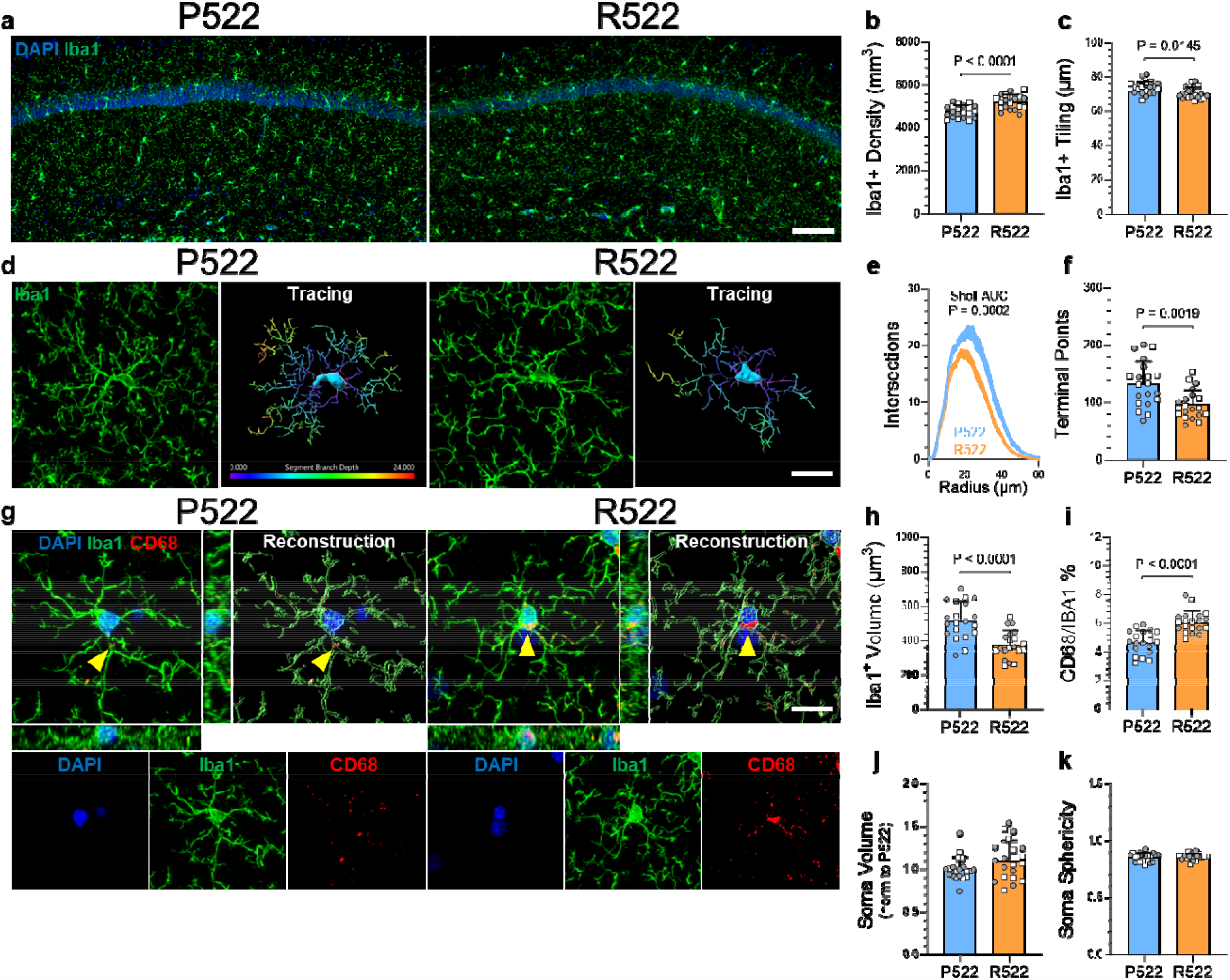
Plcg2^R522^ variant induces microglia density and morphology changes in the hippocampus of adult wildtype mice. a Representative image of microglia staining (Iba1, green) from the hippocampus (CA1) of wildtype mice (Plcg2^P522^) and Plcg2^R522^ variant. Scale bar 100 μm. b Hippocampal CA1 Iba1^+^ cell density. c Hippocampal CA1 microglia tiling score (average distance to 3 nearest Iba1^+^ microglia). d Representative 3D image of Iba1^+^ microglia morphology from the hippocampus (CA1) of wildtype mice (Plcg2^P522^) and Plcg2^R522^ variant with accompanying 3D traced reconstruction coloured according to cell soma (cyan) and branch depth. Scale bar 30 μm. e Sholl analysis of Iba1^+^ microglia. f Number of terminal points from Iba1^+^ microglia. g Representative 3D image of CD68^+^ lysosomal staining (red, yellow arrowheads) in Iba1^+^ microglia (green), with the accompanying XZ and YZ views and 3D reconstruction. Scale bar 30 μm. h Microglia cell volume. i Percentage of CD68^+^ puncta inside Iba1^+^ microglia. j-k Hippocampal microglia soma volume and soma sphericity measurements. All data points represent individual mice; N = 10 males (squares) and 10 females (circles) from both genotypes. b-c Data represented as the average of 3 entire CA1 stratum radiatum hippocampal fields viewable within 2.40 mm^2^ field of view (10x objective). e-f and h-k Data represent an average of 10 3D microglia images per mouse (d = 20x objective and g = 63x objective). Data were analysed by Two-way ANOVA considering genotype and sex; no sex differences were detected in these datasets. P values represent the effect of genotype (Plcg2 R522). b Interaction P = 0.4041, Sex P = 0.4929, Genotype P < 0.0001. c Interaction P = 0.8990, Sex P = 0.9064, Genotype P = 0.0145. e Interaction P = 0.4511, Sex P = 0.5974, Genotype P =0.0002. f Interaction P = 0.5354, Sex P = 0.6255, Genotype = 0.0019. h Interaction P = 0.6488, Sex P = 0.1702, Genotype P < 0.0001. i Interaction P = 0.7135, Sex P = 0.3667, Genotype P < 0.0001. j Interaction = 0.5483, Sex P = 0.6648, Genotype P = 0.1066. k Interaction P = 0.5381, Sex P = 0.5187, Genotype P = 0.7083.

Given the relevance of the P522R *PLCG2* variants to AD, we introduced the variant into the *App*^*NL-G-F*^ mouse model to assess how the R522 variant influences the pathology in the context of an amyloid-driven mouse model of AD. At 6 months of age, *App*^*NL-G-F*^ mice display an abundance of amyloid plaques in the brain^15^. Using Thioflavin-S (ThioS) to label fibrillar amyloid, plaque coverage was rather counterintuitively found to be increased in the hippocampus (**Fig. 2 a-b**), with more numerous individual plaque cores and larger, more densely compacted plaques (**Fig. 2 c-e**) in the *Plcg2*^R522^-expressing mice, compared to the *Plcg2*^*P522*^ mice. In the cortex (**Supp Fig. 2 a-d**), plaque burden was also influenced by sex, with female *App*^*NL-G-F*^ mice exhibiting an increased plaque burden compared to their male counterparts. The R522 variant was associated with increased plaque prevalence in the cortex of male *App*^*NL-G-F*^ mice. Despite the female plaque load remaining unchanged in the *Plcg2*^*R522*^-expressing mice, the plaques were more densely compacted than those in the *Plcg2*^*P522*^ mice (**Supp Fig. 2 e)**.

**Fig. 2:**
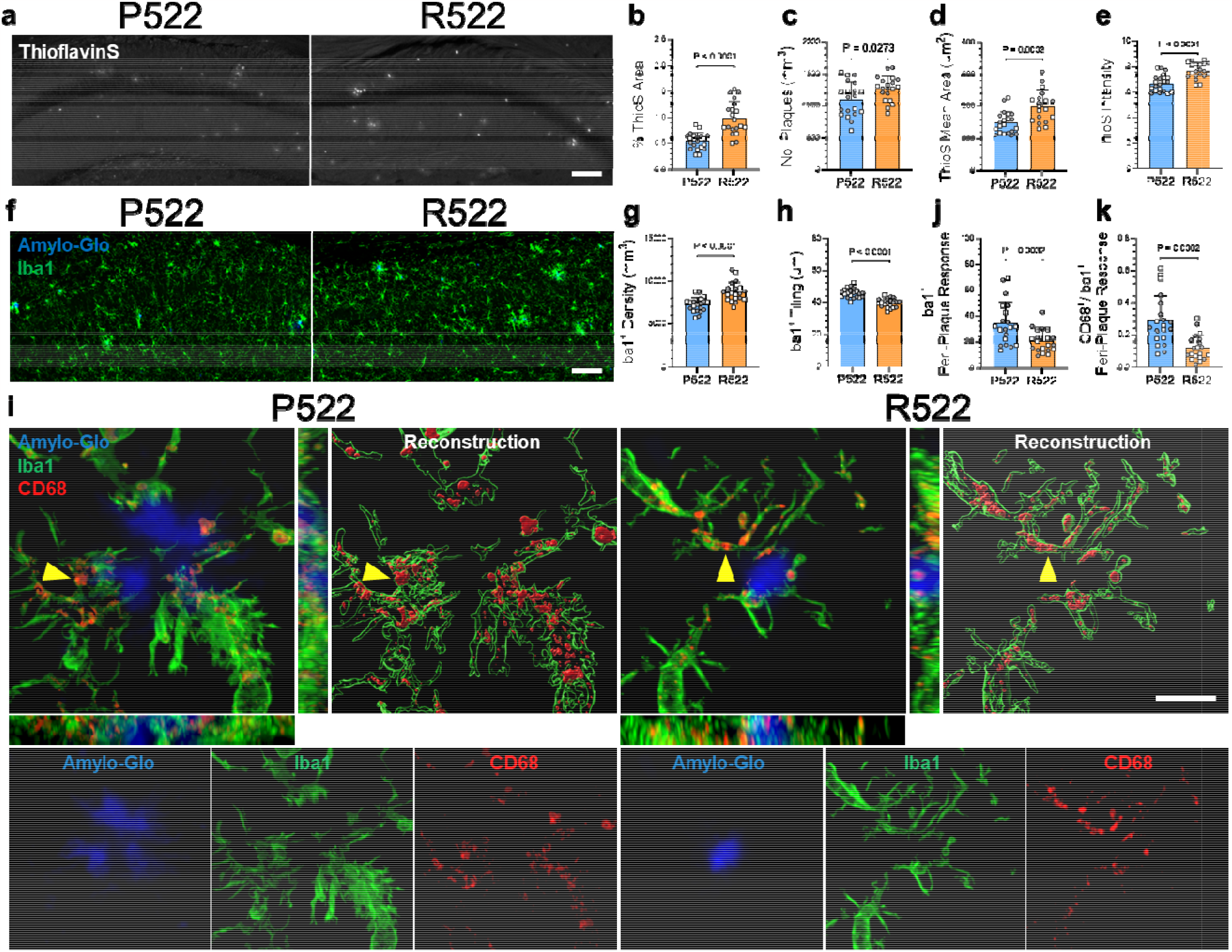
Amyloid plaque burden is increased with Plcg2^R522^ variant expression in App^NL-G-F^ mice. a Representative images of Thioflavin-S (ThioS) plaque deposition (white) from the hippocampus (CA1) of App^NL-G-F^ mice expressing either Plcg2^P522^ or Plcg2^R522^ variant. Scale bar 100 μm. b-e Hippocampal plaque parameters: ThioS plaque area coverage, Number of ThioS plaques, Individual ThioS plaque core size and ThioS plaque intensity. f Hippocampal CA1 Iba1^+^ microglia (green) co-stained with Amylo-Glo (blue). Scale bar 100 μm. g Hippocampal CA1 Iba1^+^ cell density. h Microglia tiling score (average distance to 3 nearest Iba1+ microglia). i Representative 3D image of hippocampal CA1 plaques for analysis of peri-plaque regions of interest (30 μm radius around plaque core) for Iba1^+^ microglia (green) and CD68 lysosomal marker (red, yellow arrowheads) in proximity to Amylo-Glo^+^ plaque core (blue), with the accompanying XZ and YZ views and 3D reconstruction. Scale bar 20 μm. j Hippocampal Iba1^+^ peri-plaque response. k Hippocampal CD68^+^ to Iba1^+^ peri-plaque response. All data points represent individual mice; N = 10 males (squares) and 10 females (circles) from both genotypes. b-e and g-h Data represented as the average of 3 entire CA1 stratum radiatum hippocampal fields viewable within 2.40 mm^2^ field of view (10x objective). j-k Data represent an average of 5 3D plaque core images of similar size per mouse (63x objective). Data were analysed by Two-way ANOVA considering genotype and sex; no sex differences were detected for the hippocampus datasets. P values represent the effect of genotype (Plcg2 R522). b Interaction P = 0.6898, Sex P = 0.4900, Genotype P < 0.0001. c Interaction P = 0.2533, Sex P = 0.0920, Genotype P = 0.0273. d Interaction P = 0.2497, Sex P = 0.2895, Genotype P = 0.0003. e Interaction P = 0.4922, Sex P = 0.8689, Genotype P < 0.0001. g Interaction P = 0.0873, Sex P = 0.5675, Genotype P < 0.0001. h Interaction P = 0.1508, Sex P = 0.5342, Genotype P < 0.0001. j Interaction P = 0.8787, Sex P = 0.6225, Genotype P = 0.0032. k Interaction P = 0.4066, Sex P = 0.8427, Genotype P = 0.0002.

We next examined the microglia response in the *App*^*NL-G-F*^ mice with and without the *Plcg2*^*R522*^ variant. Global scoring of microglia in the hippocampus (**Fig. 2 f-h)** and cortex (**Supp Fig. 2 f-h**) revealed an increase in microglia tiling densities, however in the cortex, the increases were only observed in the male mice. Compared to *Plcg2*^*P522*^-expressing mice, *Plcg2*^*R522*^-expressing mice demonstrated reduced peri-plaque microglia immunoreactivity (defined as being within 30 μm of a plaque) (**Fig. 2 i-j**). Additionally, lysosomal activity (CD68) was also reduced in *Plcg2*^*R522*^-expressing peri-plaque microglia compared to their *Plcg2*^*P522*^ counterparts (**Fig. 2 k**) and in contrast to the differences seen in the absence of *App*^*NL-G-F*^.

Since the R522 variant reduces the microglial and lysosomal peri plaque immunoreactivity, we next assessed whether the variant influenced synapses. In mice with the R522 variant of PLCγ2, engulfment of synaptic puncta (PSD95), detected by colocalization with microglial CD68, was lower within peri-plaque regions, whilst the surrounding peri-plaque synaptic puncta density was higher (**Fig. 3 a-b**) than in the mice expressing the P522 variant. Similar results were obtained with VGLUT2, a component of excitatory glutamatergic synapses (**Supp Fig. 3 a-b**). Given the observed *Plcg2* genotype-dependent differences in peri-plaque synaptic densities, we undertook a global measure of the synaptic puncta density in the hippocampal CA1 region assessing presynaptic bassoon and postsynaptic PSD95 markers (**Fig. 3 c-e**). *App*^*NL-G-F*^ mice, which express the risk P522 variant of PLCγ2, displayed pronounced synaptic puncta loss in both pre and postsynaptic markers, which did not occur in the *App*^*NL-G-F*^ mice with the PLCγ2 R522 variant. No changes to the density of either synaptic puncta marker were observed by the R522 variant alone in the absence of the *App*^*NL-G-F*^ transgene (**Fig. 3 c-e**).

**Fig. 3:**
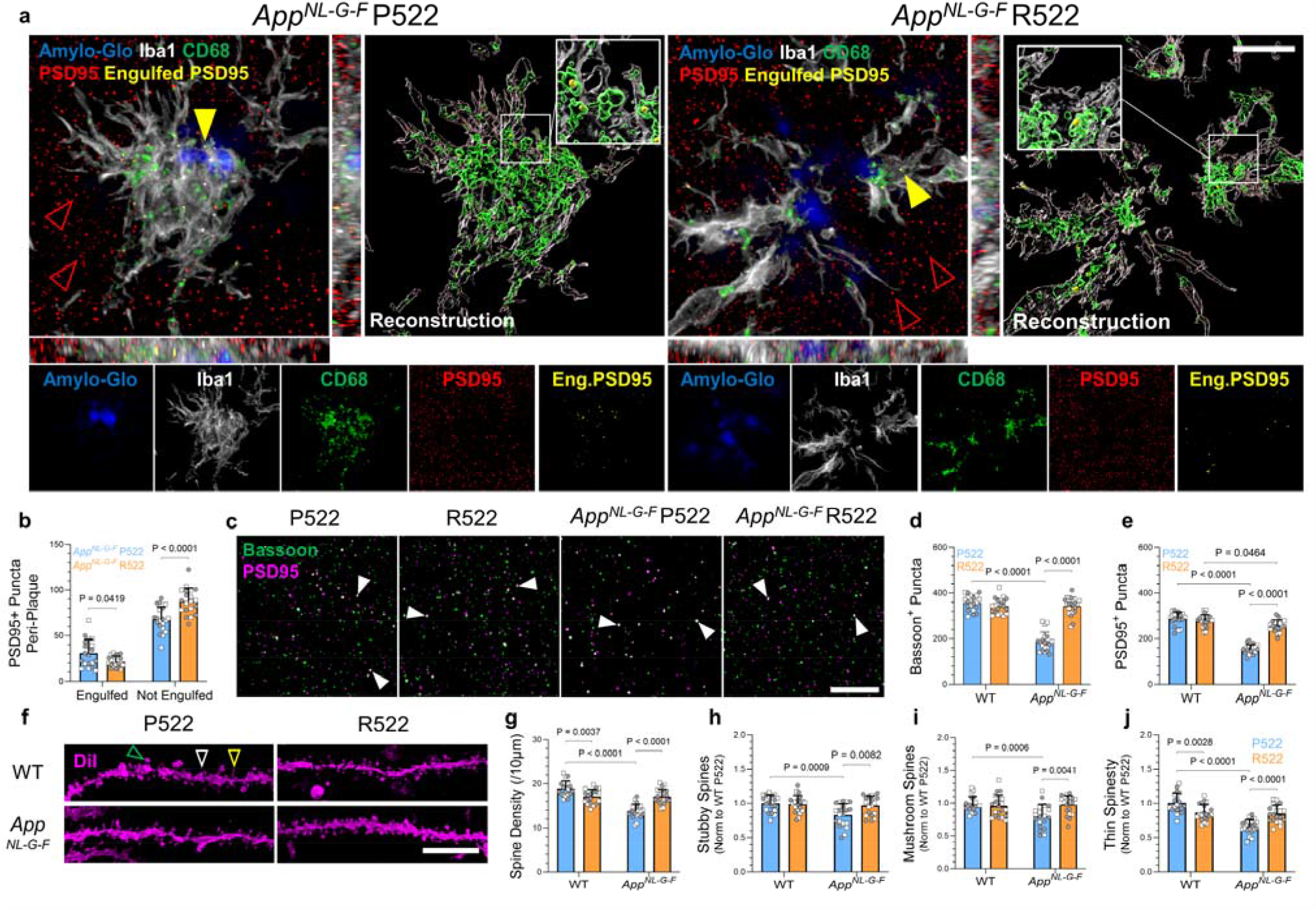
Plcg2^R522^ expression protects synapses in App^NL-G-F^ mice. a Representative 3D image of hippocampal CA1 plaques for analysis of periplaque regions of interest (within 30 μm radius around plaque core) for synaptic engulfment in App^NL-G-F^ mice with the risk P522 and protective R522 Plcg2 variants. Amylo-Glo (blue), CD68 (green), PSD95 (red, synaptic puncta) and engulfed PSD95 (yellow). Yellow arrowheads indicates areas of engulfed PSD95 inside CD68 lysosomes, empty red arrowheads indicate not engulfed PSD95. Scale bar 5 μm. b Quantification of hippocampal CA1 PSD95 synaptic puncta peri-plaque separated between engulfed (i.e. within CD68^+^ lysosomes) and not engulfed puncta surrounding the plaque. White arrowheads indicates areas pre- and postsynaptic colocalization. c Hippocampal CA1 presynaptic bassoon (green) and postsynaptic PSD95 (magenta) synaptic puncta greater than 30 μm from plaque cores from wildtype and App^NL-G-F^ mice with the risk P522 and protective R522 Plcg2 variants. Scale bar 10 μm. d-e Quantification of hippocampal CA1 bassoon and PSD95 synaptic puncta greater than 30 μm from plaque cores in WT and App^NL-G-F^ mice with the risk P522 and protective R522 Plcg2 variants. f DiOlistic labelled hippocampal CA1 dendrites with spine protrusions. Empty arrowheads correspond to stubby (white), mushroom (green) and thin (yellow) spine subtypes. Scale bar 10 μm. g Overall spine densities. h-j Relative proportions of stubby, mushroom and thin spine subtypes (based on morphological head and neck parameters). All data points represent individual mice; N = 10 males (squares) and 10 females (circles) from both genotypes. B Data represent an average of 5 plaque cores of similar sizes from the CA1 stratum radiatum hippocampal fields. D-e Data represented as the average of 6 (30 μm x 30 μm) ROI from the CA1 stratum radiatum hippocampal field per mouse. G-j Data represented as the average of 10 dendritic segments traced per mouse from the CA1 hippocampal apical secondary dendrites within the stratum radiatum. Data were analysed by Two-way ANOVA considering genotype and sex for b, and Three-way ANOVA considering Plcg2 P522R variants, App^NL-G-F^ status and sex for d-e and g-j; no sex differences were detected for the hippocampus datasets and therefore the sex cofactor was consolidated in d-e and g-j for Two-way ANOVA considering both Plcg2 and App^NL-G-F^ genotypes. All P value reported represents the post hoc multiple comparisons test (Bonferroni). B Interaction P < 0.0001, PSD95 Puncta engulfed/not engulfed P < 0.0001, Genotype P = 0.0679. d Interaction P < 0.0001, App P < 0.0001, Genotype P < 0.0001. e Interaction P < 0.0001, App P < 0.0001, Genotype P < 0.0001.g Interaction P < 0.0001, App P < 0.0001, Genotype P = 0.0369. h Interaction P = 0.0177, App P = 0.0031, Genotype P = 0.0399. i Interaction P = 0.0029, App P = 0.0137, Genotype P = 0.0753. j Interaction P < 0.0001, App P < 0.0001, Genotype P = 0.3304.

To validate the observed changes in synaptic puncta through an alternative approach, we utilised a robust method for labelling dendritic spines in the hippocampus^16,17^, which are the sites for the majority of postsynaptic sites on the CA1 neurons. *App*^*NL-G-F*^ mice displayed a prominent loss of dendritic spines compared to wildtype mice, however, the mice expressing the PLCγ2 R522 variant were protected from this (**Fig. 3 f-g**). Further investigation of dendritic spine morphology, through categorisation into stubby, mushroom and thin subtypes (**Fig. 3 h-j**), pinpointed that the *App*^*NL-G-F*^ loss of spines was evident across all subtypes and that the R522 variant conferred protection against losses of each subtype. The protective effect of the PLCγ2 R522 variant on the APP-dependent loss of dendritic spines resulted in spine densities that were comparable to those seen in similarly aged mice with the R522 variant but lacking the *App*^*NL-G-F*^ transgene. However, synaptic remodelling in wildtype mice occurred with the R522 variant, characterised by a slight decrease in the total spine number, which was associated with a reduction in thin spine subtypes compared to wildtype mice.

In summary, expression of the *Plcg2*^*R522*^ variant had a noticeable effect on microglia in adult mice. Compared to mice expressing the common P522 allele, this was characterised by an increased number of less ramified cells, which expressed higher levels of CD68. When bred onto the *App*^*NL-G-F*^ model of amyloid-driven disease, the AD-protective R522 variant of PLCγ2 resulted in an increased amyloid burden and, in spite of an increased microglial density overall, the *Plcg2*^*R522*^ expressing mice exhibited reduced microglial peri-plaque localisation. These peri-plaque microglia had a reduced CD68 expression suggestive of reduced activation and/or phagocytic activity when compared to the *Plcg2*^*P522*^ mice.

During preparation of this manuscript, Tsai et al demonstrated, using the 5xFAD mouse model, that the R522 variant of PLCγ2 resulted in reductions in both amyloid burden and its associated features of disease, compared to mice expression the P522 variant of PLCγ2^18^. The impact of PLCγ2 variants on the amyloid burden in the 5xFAD model, demonstrate the potential importance of PLCγ2 activity in regulating amyloid processing, but limited the authors ability to interpret the role of the variants in downstream pathological sequalae, such as impact on neuronal function. We utilised the knock-in *App*^*NL-G-F*^ model that benefits from physiological *App* expression levels and lacks the overexpressing artefacts found in other transgenic models^15^. Perhaps surprisingly, rather than a reduction in amyloid burden in the *App*^*NL-G-F*^ model in the presence of the R522 variant of PLCγ2, we observed an increase in amyloid load compared to the P522 variant. However, compared to the common variant, the R522 variant of PLCγ2 resulted in a striking abrogation of amyloid-induced synapse loss as measured by quantification of synaptic puncta and dendritic spines. Consistent with a direct role for microglia in this neuroprotection, the R522 variant of PLCγ2 was also associated with a significant reduction in phagocytosis of synaptic material by peri-plaque microglia.

Collectively, these observations clearly demonstrate an uncoupling of amyloid burden and synaptic loss mediated by PLCγ2. The mechanism for this is through altered microglial activity, decreased peri-plaque microglial accumulation and reduced synapse loss, associated with a drop in microglial phagocytosis of synaptic material. This novel insight, indicating that modulation of microglial activation can directly regulate synaptic clearance independently of plaque burden, consolidates the case for independent therapeutic targeting of PLCγ2 for neuroprotection and cognitive resilience in AD. Consistent with this, and demonstrating the relevance of our observations to human disease and aging, the R522 variant of PLCγ2 is in humans associated with longevity and protection of cognitive function in old age^4^ and protects against AD^1^, as well as frontotemporal dementia and dementia with Lewy bodies^4^. Furthermore, cognitive decline itself is poorly correlated with changes in amyloid burden and significant numbers of aged cognitively normal individuals exhibit some level of amyloid deposition suggesting that other mechanisms drive symptom development^19^. The first therapies for AD, targeting the removal of amyloid, are now being approved with modest positive outcomes, although associated with several side-effects^8–10^. Our data indicate that targeting PLCγ2 therapeutically, would provide an alternative strategy that could be employed independently, or as an adjunct to current amyloid-directed approaches, and may have broadly applicability in neuroinflammatory conditions beyond the scope of Alzheimer’s disease.

## Methods

### Animal Models

All experiments were approved by the Animal Welfare and Ethical Review Body (AWERB) - subgroup of the Biological Standards Committee - and conducted in accordance with UK Home Office Guidelines and Animal [Scientific Procedures] Act 1986, which encompasses EU Directive 2010/63/EU on the protection of animals used for scientific purposes. Mouse models were based on the B6.Cg-*Plcg2*^*em1Msasn*^/J (*Plcg2*^*R522*^) mice, as we previously described^13^ and available from The Jackson Laboratory (strain: 029598). Control mice were obtained from littermate founders (*Plcg2*^*P522*^). Both *Plcg2* genotypes were crossed to the knock-in *App*^*NL-G-F/NL-G-F*^ mice carrying the APP Swedish (KM670/671NL), Iberian (I716F), and Arctic (E693G) mutations to model amyloid-driven pathology^15^. Mouse genotypes were determined using real-time PCR with specific probes designed for each gene (Transnetyx, Cordova, TN). All mice were maintained as homozygous animals and co-housed in temperature-controlled, pathogen-free conditions with a 12-hour light/dark cycle and ad libitum access to food and water. All mice were aged to 6 months for the study and both male and female mice were used in all assays with sex included as a factor in all analyses.

### Brain extraction and preparation

Mice were euthanised through an intraperitoneal administration of an overdose of Euthatal® (Merial Animal Health Ltd). After confirmation of death, mice were transcardially perfused with phosphate-buffered saline (PBS) and brain tissue was extracted from the skull and fixed in 1.5% PFA (Sigma, 1.00496) for at least 72 hours (concentration dependent on downstream assays). The fixed brains were then transferred and stored in PBS containing 0.1% Sodium Azide (Sigma) at a temperature of 4°C. Subsequently, the brains were sectioned into free-floating coronal sections with a thickness of 50 μm using a Leica VT1200S vibratome (Leica Biosystems). Coronal sections ranging from -1.34 mm to -2.18 mm from the bregma were selected from each mouse, with the mouse brain atlas serving as a visual reference for analysing hippocampal and cortical regions.

### Amyloid plaque Thioflavin S Staining

Brain sections were incubated with Thioflavin-S (0.1% (w/v), Sigma, T1892) dissolved in 70% ethanol for 20 minutes at room temperature. Sections were washed in 50% ethanol three times and rehydrated in PBS prior to mounting using VECTASHIELD Vibrance Antifade Mounting Medium (Vector, H-1700) and stored in darkness at 4°C until imaging.

### Immunofluorescence

The brain sections underwent Heat-Induced Epitope Retrieval (citrate, Abcam, ab93678) for 30 minutes followed by 1% Triton X-100 permeabilization for 10 minutes. Sections were washed three times with 0.05% Tween-20 in PBS and incubated in a blocking buffer (comprising 5% species-relevant normal sera in 0.05% Tween-20 in PBS) for an hour at room temperature on an orbital shaker. The sections were then incubated with primary antibodies in blocking buffer for 48 hours at 4°C. Primary antibodies: rabbit anti-Iba1 (1:1000, FUJIFILM Wako, 019-19741), chicken anti-Iba1 (1:250, Synaptic Systems, 234009), rat anti-CD68 (1:500, Bio-Rad, MCA1957), rabbit anti-PSD95 (1:500, Abcam, ab18258), guinea pig anti-Bassoon (1:500, Synaptic Systems, 141004), guinea pig anti-VGLUT2 (1:500, Synaptic Systems, 135404). Sections were washed three times in PBS and incubated with the appropriate species-specific Alexa Fluor secondary antibodies (1:500, Invitrogen, A32740, A11006, A32759, A32740, A11073) in blocking buffer for 2 hours at room temperature on an orbital shaker. After washing in PBS, the sections were counterstained with DAPI (Invitrogen) and endogenous autofluorescence was quenched using 0.1% Sudan Black B (Sigma, 199664) in 70% ethanol, or when assaying synaptic puncta engulfment with TrueBlack Lipofusin Autofluorescence Quencher (according to manufacturer instructions, Biotium, 23007). Isotype controls (Abcam and Invitrogen) and secondary-only controls were used during the optimisation process to ensure signal specificity. Where necessary, amyloid plaques were counterstained using Amylo-Glo (according to manufacturer instructions, Biosensis, TR-300-AG). The brain slices were then mounted using VECTASHIELD Vibrance Antifade Mounting Medium (Vector, H-1700) and stored in the dark at 4°C until imaging.

### DiOlistic Spine Labelling

Free-floating brain sections underwent DiOlistic labelling to assess the hippocampal dendritic spine densities and spine morphologies. DiOlistics was performed as previously described^16,17,20^. Briefly, on glass slides 1 μm tungsten particles (40 mg, Fisher, 11319718) were coated dropwise with 1,1′-Dioctadecyl-3,3,3′,3′-Tetramethylindocarbocyanine Perchlorate (DiI, 2 mg, Invitrogen, D282) dissolved in dichloromethane (400 ul, Sigma, L090001). DiI-coated tungsten microcarriers were transferred to Tefzel tubing (Bio-Rad, 1652441) and cut into ‘bullets’ for delivery using Helios Gene Gun System (Bio-Rad, 1652432). Brain sections were positioned on histology slides, and bullets were fired at 80-100 psi through an inverted cell culture insert (8.0 μm, Sigma, BR782746-24EA). Sections were immersed in PBS for 1 hour at room temperature for sufficient dye diffusion. Sections were fixed in 4% PFA (Sigma, 1.00496) for 10 minutes to halt dye diffusion, washed in PBS, and nuclei stained with DAPI. DiOlistic labelled tissue sections were mounted in glycerol-free mounting reagent FluorSave (Millipore, 345789). Sections were stored in the dark, and due to the nature of DiI labelling were imaged the same day.

### Immunofluorescence and Thioflavin S Image Acquisition

Imaging was performed using a Lecia SP8 Lightning confocal microscope with Leica HyD photodetectors set with a minimum 10% gain and fixed laser powers for each experiment. Iba1^+^ cell bodies and Thioflavin S^+^ plaque coverage were imaged with 10x objective, scan speed of 600Hz, line average of 3 covering a field of view size of 1,500 μm x 1,500 μm (voxel size of 0.379 μm) through a depth of 60 μm with intervals of 5 μm. Images were compressed to a 2D maximum projection for analysis. Images of microglia morphology were captured using a 20x objective, scan speed of 400Hz, and line average of 3 covering a field of view size of 775 μm x 775 μm (voxel size of 0.189 μm) through a depth of 40 μm with images analysed as 3D stacks. Punctate staining, CD68 and synaptic puncta were captured using 63x objective, scan speed 600Hz, line average of 3 covering a field of view size of 246 μm x 246 μm (voxel size of 0.06 μm) through a depth of 5 μm. For synaptic puncta staining alone, only a depth of 1 μm from the initial 5 μm was analysed, for CD68 (including containing with synaptic puncta) the entire 5 μm stack was analysed. Images were pseudo-coloured for visualisation of immunostaining.

### DiOlistic Spine Image Acquisition

Dendritic spines labelled by DiOlistics were imaged from CA1 secondary dendrites within the striatum radium of the hippocampus and were selected based on minimal overlap with adjacent cells. The Lecia SP8 Lightning confocal microscope was configured under the resonant scanner with PMT photodetector set with a gain of 400 and fixed laser powers. Spines were captured using 63x objective, scan speed of 8000Hz, line average of 2 covering a field of view size of 147.72 μm x 147.72 μm (voxel size of 0.144 μm) through a varying z depth typically spanning 20-50 μm dependent of labelling. Images were subsequently deconvolved using Leica Lightning Deconvolution software to resolve spine morphologies.

### Immunofluorescence Image Analysis

All images were analysed using Imaris (v10.0, Bitplane) and batch analysed through a sequential combination of the images into a ‘time-series’ for each experimental marker. For 10x objective images, ROIs of the entire hippocampal CA1 striatum radium field and cortex were extracted using manual Surface drawing. For 20x objective microglia morphology images and 63x objective CD68 puncta in microglia, circle ROI microglia territory fields (50 μm radius) were placed over cells with suitable single-cell territories and extracted from the original images. For 63x objective synaptic puncta staining, 6 smaller ROI fields 30 μm x 30 μm (excluding cell nuclei and greater than 30 μm from plaques where relevant) were positioned and extracted from the original images. For 63x peri-plaque analysis, circle ROI territory fields (30 μm radius) were extracted over plaque cores and extracted from original images. In all analysis pipelines the fluorescence intensities across the extracted regions were normalised across the batch ‘time-series’ using the Imaris XTension ‘Normalise Time Points’ and applied with a Gaussian filter. For 10x objective images, Iba1^+^ cell bodies were quantified using the Spot function and Thioflavin-S plaques were identified using the Surface function. For 20x objective images, microglia morphology was quantified using the Surface function to ‘isolate’ single microglia based on the largest surface within the circle ROI territory fields. Single microglia surfaces were then extracted and morphologically assessed using the Filament Tracer function, with AI segment features based on the seed point of 0.2 μm. For 63x objective images of microglia with CD68 puncta were analysed using the Surface function to isolate the microglia channel within the circle ROI territory fields and masked onto the CD68 channel, with the CD68 puncta quantified with the Surface function. For 63x objective images for synaptic puncta, punctate staining was analysed using the Surface function. For 63x objective images for peri-plaque analysis, the Surface function was used to isolate the plaque core and microglia immunoreactivity, with the microglia surface render masked onto the CD68 channel and, where relevant, the CD68 channel masked onto the synaptic puncta channel for assessing synaptic engulfment. Synaptic puncta staining was surface rendered for the original puncta staining channel and the masked engulfed synaptic puncta that colocalise with CD68 within microglia. An animation of the engulfed synaptic puncta periplaque in *App*^*NL-G-F*^ mice is demonstrated in **Supp Fig 4**.

### DiOlistic Spine Image Analysis

Dendritic spines were analysed using the Filament Tracer function in Imaris (v9.9, Bitplane). ‘ROIs’, non-overlapping areas free from dye debris and other labelling, were positioned along dendrites greater than 30 μm in length. The dendrite base shaft was traced using default thresholding followed by spine tracings using fixed parameters of spine head seed point size of 0.45 μm and manually checked to ensure correct tracing. Spines were automatically subclassified (stubby, mushroom, thin) using SpineClassifier MATLAB extension, distinguishing based on spine length and head size with the following fixed parameters: Stubby = length(spine) < 0.8 [short protrusions without presence of spine neck], Mushroom = length(spine) < 3 and max_width(head) > mean_width(neck) [spine protrusions with defined and obvious spine head], Filopodia = length(spine) > 3 [this spine subtype was removed from analysis], and Thin = true [all remaining spines].

### Statistics

All data analysis was collected blinded to genotype and sex, with individual experiments (i.e. one experimental marker/readout) analysed in one batch by a single operator. All graphs and statistical analyses were generated using GraphPad Prism (GraphPad Software, version 10). The Shapiro–Wilk test was used to check for normal distribution in all datasets. Data was analysed by Three-way and Two-way ANOVA tests for genotypes and sex and, where appropriate post hoc multiple comparisons (Bonferroni). Data is presented as an average value for individual mice generated from N = 10 males and 10 females from all genotypes with mean ± standard deviation (SD), apart from Sholl analysis which is presented as mean ± standard error of the mean (SEM). All P values are listed either in the necessary figure or in the accompanying figure legends.

## Acknowledgements

This work is supported by the UK Dementia Research Institute [award number UK DRI-3001], which receives its funding from UK DRI Ltd, funded by the UK Medical Research Council, Alzheimer’s Society and Alzheimer’s Research UK. P.R.T is, in part, funded by The Moondance Foundation.

## Author Contributions

R.J.B performed experiments and analysed data. R.J.B, E.M, T.P, E.S and M.V provided methodology. R.J.B, E.M, T.R, E.S, M.V, J.W and P.R.T provided intellectual contributions. R.J.B and P.R.T wrote the manuscript. P.R.T conceived the study. All authors reviewed the manuscript.

