## Supplementary material for "Uncoupling of synaptic loss from amyloid burden by an Alzheimer’s disease protective variant of PLCγ2": Supp Fig

**Author affiliations:**
[1] UK Dementia Research Institute at Cardiff, Cardiff University, Cardiff, CF24 4HQ, UK.
[2] Systems Immunity Research Institute, Heath Park, Cardiff University, Cardiff, CF14 0NN, UK.

Correspondence to: Professor Philip R. Taylor
UK Dementia Research Institute at Cardiff, Hadyn Ellis Building, Maindy Road, Cardiff University, Cardiff, CF24 4HQ, UK;


### Supplementary Data

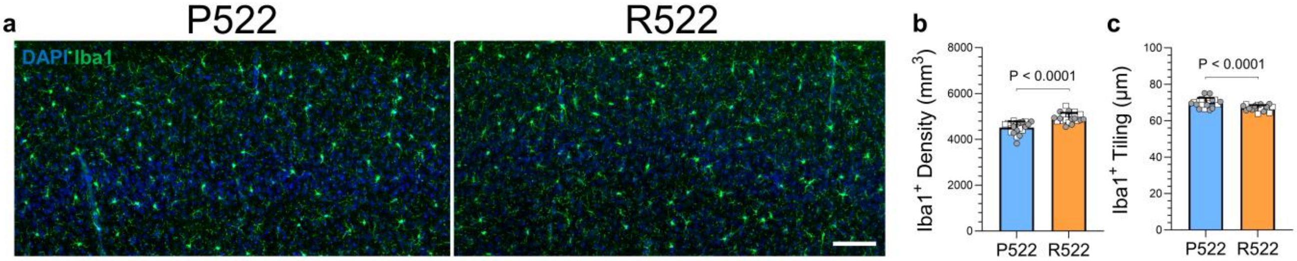


***Supp Fig. 1: Plcg2$^{R522}$ variant increases microglia density in the cortex of adult wildtype mice.*** *a Representative image of microglia staining (Iba1, green) from the cortex of wildtype mice (Plcg2$^{P522}$) and Plcg2$^{R522}$ variant. Scale bar 100 µm. b Cortex Iba1$^{+}$ cell density. c Cortex microglia tiling score (average distance to 3 nearest Iba1$^{+}$ microglia). All data points represent individual mice; N = 10 males (squares) and 10 females (circles) from both genotypes. Data represented as the average of 3 cortex fields viewable within 2.40 mm$^{2}$ field of view (10x objective). Data were analysed by Two-way ANOVA considering genotype and sex; no sex differences were detected in these datasets. P values represent the effect of genotype (Plcg2 R522). b Interaction P = 0.9367, Sex P = 0.2798, Genotype P < 0.0001. c Interaction P = 0.5629, Sex P = 0.9581, Genotype P < 0.0001.*

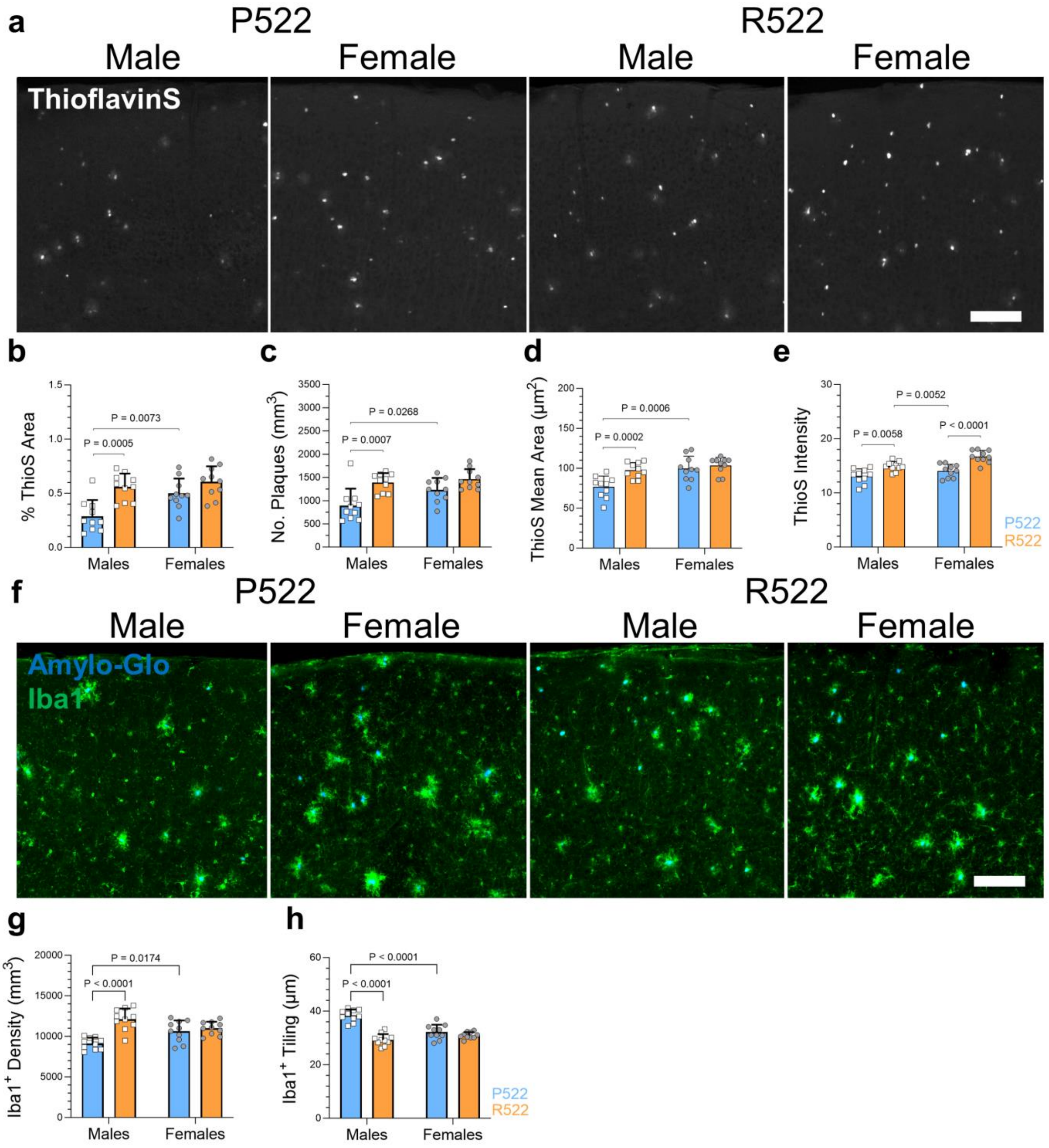


***Supp Fig. 2 Plcg2$^{R522}$ variant increases male cortical amyloid plaque burden in App$^{NL\text{-}G\text{-}F}$ mice.*** *a Representative images of Thioflavin-S (ThioS) plaque deposition (white) from the cortex of App$^{NL\text{-}G\text{-}F}$ mice expressing either Plcg2$^{P522}$ or Plcg2$^{R522}$ variant. Scale bar 100 µm. b-e Cortex plaque parameters: ThioS plaque area coverage, Number of ThioS plaques, Individual ThioS plaque core size and ThioS plaque intensity. f Cortex Iba1$^{+}$ microglia (green) coverage co-stained with Amylo-Glo (blue). Scale bar 100 µm. g Cortex Iba1$^{+}$ cell density and h Microglia tiling score (average distance to 3 nearest Iba1+ microglia). All data points represent individual mice; N = 10 males (squares) and 10 females (circles) from both genotypes. Data represent an average of 3 cortex fields viewable within 2.40mm$^{2}$ field of view (10x objective). Data were analysed by Two-way ANOVA considering genotype and sex; sex differences were detected for the cortex datasets. All P value reported represents the post hoc multiple comparisons test (Bonferroni). b Interaction P = 0.0724, Sex P = 0.0062, Genotype P = 0.0001. c Interaction P = 0.1178, Sex P = 0.0186, Genotype P < 0.0001. d Interaction P = 0.0353, Sex P = 0.0007, Genotype P = 0.0032. e Interaction P = 0.2649, Sex P = 0.0005, Genotype P < 0.0001. g Interaction P = 0.0005, Sex P = 0.5755, Genotype P < 0.0001. h Interaction P < 0.0001, Sex P = 0.0022, Genotype P < 0.0001.*

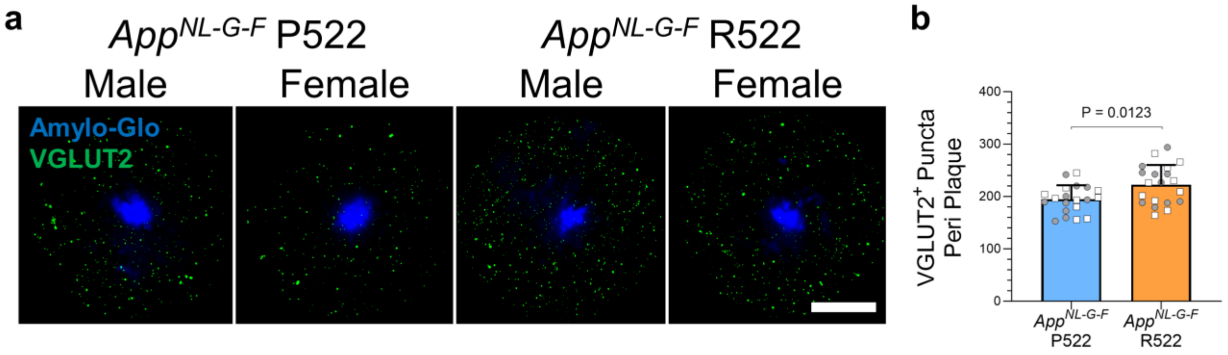


***Supp Fig. 3 $Plcg2^{R522}$ expression protects VGLUT2 synapses in $App^{NL\text{-}G\text{-}F}$ mice.*** *a Representative peri-plaque regions of interest (within 30 µm radius around plaque core) from $App^{NL\text{-}G\text{-}F}$ mice with the risk P522 and protective R522 Plcg2 variants stained for Amylo-Glo (blue) and VGLUT2 synaptic puncta (green). Scale bar 5 µm. b Quantification of VGLUT2 synaptic puncta peri-plaque. All data points represent individual mice; N = 10 males (squares) and 10 females (circles) from both genotypes. b Data represented as the average of 5 plaque cores of similar sizes from the CA1 stratum radiatum hippocampal fields. Data were analysed by Two-way ANOVA considering genotype and sex; no sex differences were detected for the hippocampus datasets. P value reported represents the Two-way ANOVA effect of genotype (Plcg2 P522R variants). b Interaction P = 0.7130, Sex P = 0.8751, Genotype P =* 0.0123.

See file “Supp Figure 4 MP4”

***Supp Fig. 4: Imaris rendered animation of peri-plaque engulfment of synaptic puncta in AppNL-G-F mice. Amylo-glo (blue), Iba1 (white), CD68 (green), PSD95 (red) and engulfed PSD95 (yellow). Scale bar 5 µm***
